# CD47-SIRPα controls ADCC killing of primary T cells by PMN through a combination of trogocytosis and NADPH oxidase activation

**DOI:** 10.1101/2022.03.08.483323

**Authors:** Françoise Gondois-Rey, Thomas W. Miller, Jacques A. Nunès, Daniel Olive

## Abstract

Immunotherapies targeting the “don’t eat me” myeloid checkpoint constituted by CD47 SIRPα interaction have promising clinical potential but are limited by toxicities associated with the destruction of non-tumor cells. These dose-limiting toxicities demonstrate the need to highlight the mechanisms of anti-CD47-SIRPα therapy effects on non-tumor CD47-bearing cells. Given the increased incidence of lymphopenia in patients receiving anti-CD47 antibodies, and the strong ADCC effector function of Poly Morpho Nuclear Cells (*PMNs*), we investigated the behavior of primary PMNs cocultured with primary T cells in the presence of anti-CD47 mAbs. PMNs killed T cells in a CD47-mAb-dependent manner and at a remarkably potent PMN to T cell ratio of 1:1. The observed cytotoxicity was produced by a novel combination of both trogocytosis and a strong respiratory burst induced by classical ADCC and CD47-SIRPα checkpoint blockade. The complex effect of the CD47 blocking mAb could be recapitulated by combining its individual mechanistic elements: ADCC, SIRPα blockade and ROS induction. Although previous studies had concluded that disruption of SIRPα signaling in PMNs was limited to trogocytosis-specific cytotoxicity, our results suggest that SIRPα also tightly controls activation of NADPH oxidase, a function demonstrated during differentiation of immature PMNs but not so far in mature PMNs. Together, our results highlight the need to integrate PMNs in the development of molecules targeting the CD47-SIRPα immune checkpoint and to design agents able to enhance myeloid cells function while limiting adverse effects to healthy cells able to participate in the anti-tumor immune response.

**Synopsis:** Dose-limiting toxicities demonstrate the need to investigate anti-CD47-SIRPα therapy effects on non-tumor CD47-bearing cells. We demonstrate that anti-CD47 mAbs stimulate potent killing of T cells by PMN through a novel combination of trogocytosis and ROS regulated by SIRPα.

**Graphical Abstract:** 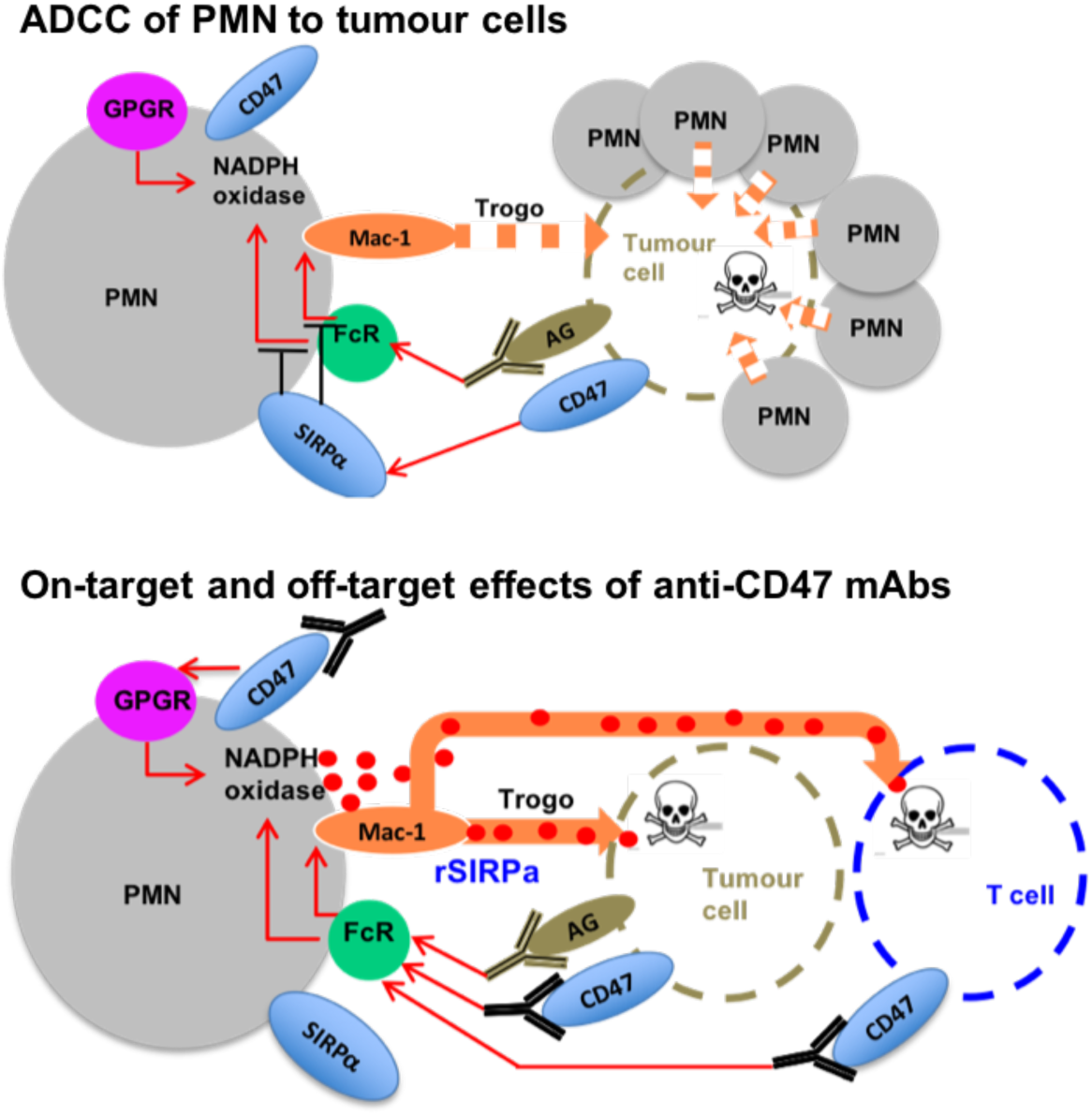

## Introduction

The ability to manipulate patients’ immunity with antibodies has changed the therapeutic outlook for cancer. Strategies based on the induction of Antibody-Dependent-Cell-Cytotoxicity (ADCC) with antibodies targeting tumor antigens like Rituximab in B-lymphomas and Trastuzumab in breast cancer have demonstrated clinical efficacy. More recently, strategies based on the blockade of the T cells inhibitory checkpoints CTLA-4 and PD-1/PD-L1 with mAbs showed spectacular efficacy but limited to certain tumor types with high mutational burdens and T cell infiltration and associated side effects (1). To create therapies with broader efficacy, strategies targeting the myeloid checkpoint, signal regulatory protein α (SIRPα) were developed to enhance myeloid ADCC induced by therapeutic antibodies (2, 3).

SIRPα controls phagocytosis when engaged by its ligand CD47, a molecule widely expressed on most cell types (4). This interaction constitutes the “don’t eat me” regulatory axis. Inhibition results in part from the Immunoreceptor Tyrosine-based Inhibitory Motif (ITIM) phosphorylation of SIRPα cytoplasmic tail that prevent modifications of the membrane leading to the formation of the phagocytic cup (5). The regulation of SIRPα signaling in PMN is known to be sensitive to their activation where IL-17 stimulation results in cleavage of the ITIM signaling domain of SIRPα (6). SIRPα also inhibits activation of Mac-1, the integrin required for spread and adhesion of myeloid cells on their target (7, 8). Mac-1 is a heterodimer of CD11b (αM) and CD18 (β2) integrins but only CD18 is required in ADCC-mediated adhesion (9). While macrophages are capable of whole-cell phagocytosis, Poly Morpho Nuclear Cells (PMNs) ingest parts of the target cell in a mechanism called trogocytosis (10, 11). In ADCC, trogocytosis is sufficient to induce necrotic cell death resulting from lytic processes but requires high ratios of PMNs per target cell (10, 12).

PMNs are mainly known for their capacity to degranulate toxic molecules accumulated during their maturation or produce toxic Reactive Oxygen Species (*ROS*) upon the “respiratory burst” within minutes of stimulation to fulfil their killing mission (13). The NADPH oxidase is a multi-subunit enzyme complex that, once assembled at the PMN membrane, generates superoxide as a precursor for others ROS (H_2_O_2_, HOCl), released in the milieu as an antimicrobial agent. However, ROS are also toxic to host cells and this process must be tightly controlled. NADPH oxidase is activated through membrane G Protein Coupled Receptors (*GPCR*) that sense the various molecules of the milieu, such as bacterial compounds (fMLP, LPS) (14), but little is known about its control. Recently SIRPα was shown to be involved in the control of NADPH oxidase by inhibiting the expression of the gp91^Phox^ subunit, a membrane component of NADPH oxidase complex, in immature cells (15). This inhibition required engagement of SIRPα by CD47 and signaling by the cytoplasmic tail of SIRPα. Blockade of SIRPα engagement resulted in enhanced production of ROS. However, NADPH oxidase and ROS are not known to be involved in ADCC (10).

CD47 is a widely expressed signaling receptor and marker of “self” involved in many biological processes through its interaction with its ligand thrombospondin-1 (*TSP-1*), an inflammatory protein that promotes migration and activation of cells (16). Different epitopes of CD47 are involved in the interaction of CD47 with TSP-1 and SIRPα (17). CD47 interacts also with SIRPγ, a molecule only expressed on T cells (18). CD47 is known to signal through its lateral association with integrins and GPCR (19). For example, triggering of CD47 induces endothelial cell spreading on RGD sequences through the lateral association of CD47 with β3 integrins and adhesion of T cells on LDV sequences through the lateral association with β1 integrins (20). An interaction of CD47 with Mac-1 was recently described as one of the mechanisms involved in fusion of macrophages (21).

The overexpression of CD47 on tumor cells suggested that blockade of the “don’t eat me” checkpoint could synergize with therapeutic mAbs (16) to enhance the elimination of tumor by myeloid cells (22). Antibodies blocking the CD47-SIRPα interaction increased phagocytosis of macrophages (22, 23) and cytotoxicity of PMNs (12), inhibited tumor engraftment (22) and eliminated pre-existing tumor in mice (24). Although both can block the “don’t eat me” interaction, anti-CD47 mAbs were more efficient than anti-SIRPα mAbs (24). This higher efficiency was thought to result from the additional ADCC resulting from opsonization of target with anti-CD47 mAbs (4, 12).

Despite their promising pre-clinical results, the clinical progress of anti-CD47 mAb therapies have been limited by on-target, non-tumor toxicities including anemia, neutropenia, thrombocytopenia and lymphopenia (25). As we had previously identified the CD47-SIRPα immune checkpoint and SIRPα activity as a key determinant of low density PMN cytotoxicity toward healthy T cells (26) we investigated the effect of the SIRPα-blocking anti-CD47 mAb (clone CC2C6) on PMN-mediated T cell cytotoxicity. By investigating the behavior of PMNs to primary T cells, we found that blockade of the engagement of SIRPα resulted in an important cytotoxicity sustained not only by an enhancement of trogocytosis but also by induction of a strong respiratory burst, resulting in suppression of T cells.

## Materials and Methods

### Cells

Blood samples were obtained from healthy donors (EFS, Etablissement Français du Sang, Marseille, France). High-density PMNs were separated from PBMC by centrifugation on Ficoll-Hypaque gradients. Red cells were eliminated with RBC lysing buffer (eBioscience, ThermoFischer, France). PMNs were kept at 4°C in PBS supplemented with Ca^2+^ 1mM and Mg^2+^ 1mM. Purity of the PMNs preparations were routinely of 70-90%, contaminants were T cells, monocytes were absent (Fig S1-a). T cells were separated from frozen or fresh PBMC using CD3^+^ magnetic beads (Miltenyi Biotech, Germany) purity of preparations was above 95% (not shown). Raji B cell lines were obtained from ATCC. For Jurkat T cell line, JA16 was initially subcloned in the lab (27) and JINB8 is a CD47deficient Jurkat cell line (28). Cells were cultivated in RPMI 1640 medium supplemented with 10% foetal calf serum and antibiotics.

### Antibodies and peptides

Purified anti-CD47 mAbs (clones CC2C6 and 2D3), purified anti-SIRPα mAbs (clone SE5A5) and G1 isotype mAbs were used at 10 µg/mL (BioLegend). Anti-Mac-1 was reconstituted by mixing anti-CD11b (clone ICRF44) and CD18 (clone TS1/18) mAbs at a final concentration of 10 µg/mL for each (BioLegend). Anti-CD3 mAbs (clone UCHT1) were prepared in the laboratory and used at 10 µg/mL. Recombinant SIRPα was prepared as in (29) and was used as a monomer or multimerized with Neutravidin at a saturating concentration of 5µM. RGDS and LGDP were used respectively at 40 µg/mL and 100 µg/mL (Sigma Aldrich Merck). 4N1K, a cell-binding domain adhesive peptide was used at 10 µM (Eurogentec).

### Cytotoxicity assay

Targets were stained with CellTrace™ Violet (Life Technologies, France). Counting beads (BD Biosciences, France) were added to target cell suspensions. Cocultures were set using PMNs and T cells from different donors at ratios ranging from 1/1 to 5/1 PMNs to target and incubated overnight at 37°C in culture medium. An aliquot of the coculture was stained and analyzed by flow cytometry immediately after mixing to verify ratios and after overnight incubation to evaluate cytotoxicity by counting live targets. Target counts were standardized to counting beads and either compared to counts of control targets cultured overnight without PMNs or to control coculture with PMNs. T cells did not proliferate during overnight culture, Raji cells slightly proliferate, Jurkat and JINB8 proliferated of a factor of 3-5. PMNs remained alive during overnight culture but showed a trend to display an activated phenotype characterized by decrease of SSC and increase of FSC and depending on the stimulation counts importantly decreased (shown in Fig S2-b). Inhibition of respiratory burst was performed by adding 50 µg/mL of catalase to digest H_2_O_2_ and 10 µM of Diphenyleneiodonium chloride (DPI) to inhibit NADPH oxidase (Sigma-Aldrich, Merck) during the overnight coculture. DPI could not be used with cell lines as target because it inhibited cells proliferation. Since cell lines proliferated during the overnight incubation cytotoxicity could no more be quantified as it was based on absolute counting of cells.

### Flow cytometry

Cell suspensions were stained with Near-IR LIVE/DEAD™ (Life Technologies), CD3-PC5 or CD19-PC5 and CD11b-PE when indicated (BD Biosciences). Samples were acquired on a FORTESSA cytometer (BD Biosciences). Data were exported and analyzed with FlowJo (RRID:SCR_008520; version 9-2, MacOS X). Counting beads and cells were gated on FSC-A/SSC-A (shown in Figure 1-d). Doublets were excluded on FSC-A/FCS-H. Dead cells were excluded on expression of the viability dye (shown in Fig S1-a). PMNs were gated as SSC^hi^ cells and CD11b expression. For analysis of trogocytosis of PMNs, targets T cells were excluded on expression of CD3. For cytotoxicity assays, targets (T cells, Jurkat T cell-lines, and Raji B lymphoma cell-line) were gated on SSC^low^FSC^hi^, cell-trace and CD3 or CD19 expression.

**Figure 1.**
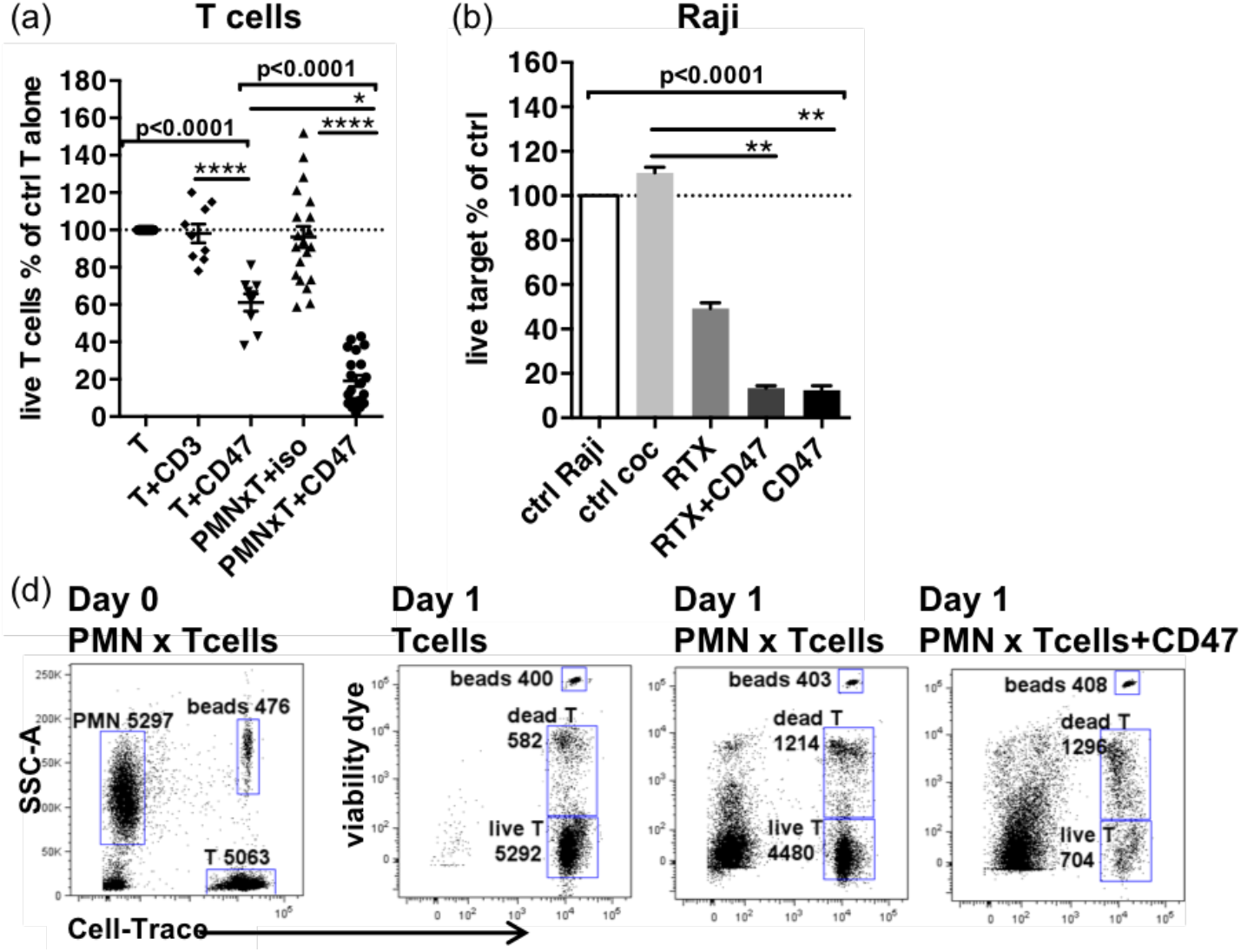
Anti-CD47 mAbs induce killing of primary T cells by PMNs. (a) Cytotoxicity to primary T cells induced by anti-CD47 mAb clone CC2C6 alone or in the presence of PMNs (n=9-25 different donors; ratio of PMN to T cells=2). Iso: isotype; CD3: anti-CD3 mAbs; CD47: anti-CD47 mAbs. (b) Induction of PMNs’ cytotoxicity to Raji lymphoma B cells by combinations of Rituximab (RTX) *plus* anti-CD47 mAb clone CC2C6 (CD47) (n=3; ratio of PMN to target=3). For (a) and (b), cytotoxicity is represented by the % of live targets in indicated conditions compared to targets cultivated overnight alone. Median and SEM. *P*-values from Kruskall-Wallis test is indicated on top of groups, *P*-values from Dunn’s multiple comparison post-test on top of pairs: *, *P*<0.05; **, *P*<0.01; ****, *P*<0.0001. (d) Cytotoxicity assay. Left plot shows coculture of PMN with cell-trace stained T cells on day 0. The following plots show cell-trace and viability staining of indicated co-cultures after overnight incubation. Gates and counts of beads, live and dead CellTrace stained T cells are shown.

### Trogocytosis

Target cells were stained with the membrane-dye PKH67 (Sigma-Aldrich) and cocultured with PMNs at ratios ranging from 1:1 to 3:1 for 3-hours. Trogocytosis was also analyzed in PMNs recovered from cytotoxicity assays after an overnight incubation with targets stained with CellTrace. Percentages of trogocytosis were determined by the expression of the T cell dye in PMNs after setting gates on PMNs cultured alone.

### Intracellular production of ROS

PMN’s were stained with 25 µM Dihydrorhodamine 123 (DHR, Sigma-Aldrich) and cultured for 1-1.5 hour in the presence of catalase at 50 µg/mL (30). Lipopolysaccharide (LPS, 100 ng/mL) *plus* N-Formylmethionyl-leucyl-phenylalanine (fMLP, 5 µM) and monoclonal antibodies at 10 µg/mL were added immediately after DHR staining. Samples were acquired on a FORTESSA cytometer, DHR was analyzed in the 525/50 channel and percentage determined by setting the gate on un-stimulated stained cells.

### Statistics

Statistical graphics were performed with Prism 6 (RRID:SCR_005375) software. Mann-Whitney, Wilcoxon-paired-matched non-parametric test or Kruskall-Wallis test followed by multiple comparison Dunn’s post-test to compare variables between groups were used as indicated.

## Data Availability

The data generated in this study are available within the article and its supplementary data files.

## Results

### Anti-CD47 mAbs induce killing of primary T cells by PMNs

To investigate the potential for CD47-dependent PMN killing of primary T cells, we treated co-cultures with anti-CD47 mAb CC2C6 at a PMN to T cell ratio of 2:1. This leads to a mean survival of 19% of control T cells after overnight coculture (Figure 1-a, Figure S2-b). Although overnight coculture with PMNs was not neutral, resulting in a variability of 50%-150% of T cells survival, and CC2C6 had a direct weak cytotoxicity to T cells previously shown (31), the effect of CC2C6 in the presence of PMNs was far more important and significant. Percentages of dead T cells in co-culture with PMNs and anti-CD47 mAbs were increased as compared to control T cells cultivated alone or with PMNs without anti-CD47 mAbs (Figure 1-d) (67% vs 15% and 22% resp.). However, the most striking difference was the important decrease of total cells suggesting that cytotoxicity manifested as necrosis rather than apoptosis, as reported for PMNs induced killing (10).

The potent CD47 mAb-induced PMN cytotoxicity was not limited to primary T cells as we observed similar cytotoxicity with Raji tumor cells (Raji) as targets. At a ratio of PMN to target of 3:1, RTX induced the killing of 50% of the Raji tumor cells (Raji) (Figure 1-b, Figure S2-a). Addition of anti-CD47 mAbs to RTX reduced the survival of Raji to 14% of control, but anti-CD47 mAbs alone reduced cell survival to 13%. The similarity of cytotoxicity induced by anti-CD47 mAbs whether alone or with RTX suggested that they were mainly responsible for the target cell death when used in the combination.

Due to the uniquely potent nature of the cytotoxicity observed for PMN in the presence of anti-CD47 antibody, we sought to investigate its mechanism.

### Trogocytosis *plus* blockade of SIRPα are insufficient to explain the enhanced cytotoxicity of PMNs in anti-CD47 mAb-triggered ADCC

Anti-CD47 mAbs are capable of exerting cytotoxic functions via a variety of mechanisms. They can simultaneously opsonize T cells to activate PMNs’ FcR providing “eat me” signals while also inhibiting “don’t eat me” signals resulting from the engagement of PMNs’ SIRPα by CD47 on T cells. PMN-mediated toxicity via ADCC was recently shown to proceed almost exclusively by trogocytosis of the target cell membrane and to be regulated by SIRPα (10). We reconstituted these mechanisms with anti-CD3 mAbs to opsonize T cells and anti-SIRPα mAbs to block “don’t eat” signals while comparing trogocytosis by PMNs and cytotoxicity to T cells.

T cells were stained by PKH67 a lipophilic dye that accumulates in the plasma membrane and whose transfer to non-labeled cells indicates trogocytosis (32). Transfer of the membrane-dye to PMNs was analyzed by flow cytometry after a 3-hours incubation. Similar percentages of PKH67^+^PMNs were found with anti-CD47 and anti-CD3 mAb treatment (59% and 60% resp., Figure 2-a) suggesting an equivalent interaction of PMNs with T cells whether opsonized with anti-CD3 or anti-CD47 mAbs. To control for non-specific cellular adhesion or ADCC mediated adhesion, we included an anti-Mac-1 mAb that would block specific ADCC mediated adhesion via integrins α_M_/β_2_. Anti-Mac-1 mAbs inhibited PMNs trogocytosis induced by anti-CD47 mAbs. Given the lateral interactions of CD47 with integrins (17), we verified whether b1 and β3 (as control) integrins were involved, but only antibodies to the b2 integrin CD18 inhibited anti-CD47 mAbs-induced trogocytosis, as reported for PMNs’ ADCC (Figure S3-b).

**Figure 2.**
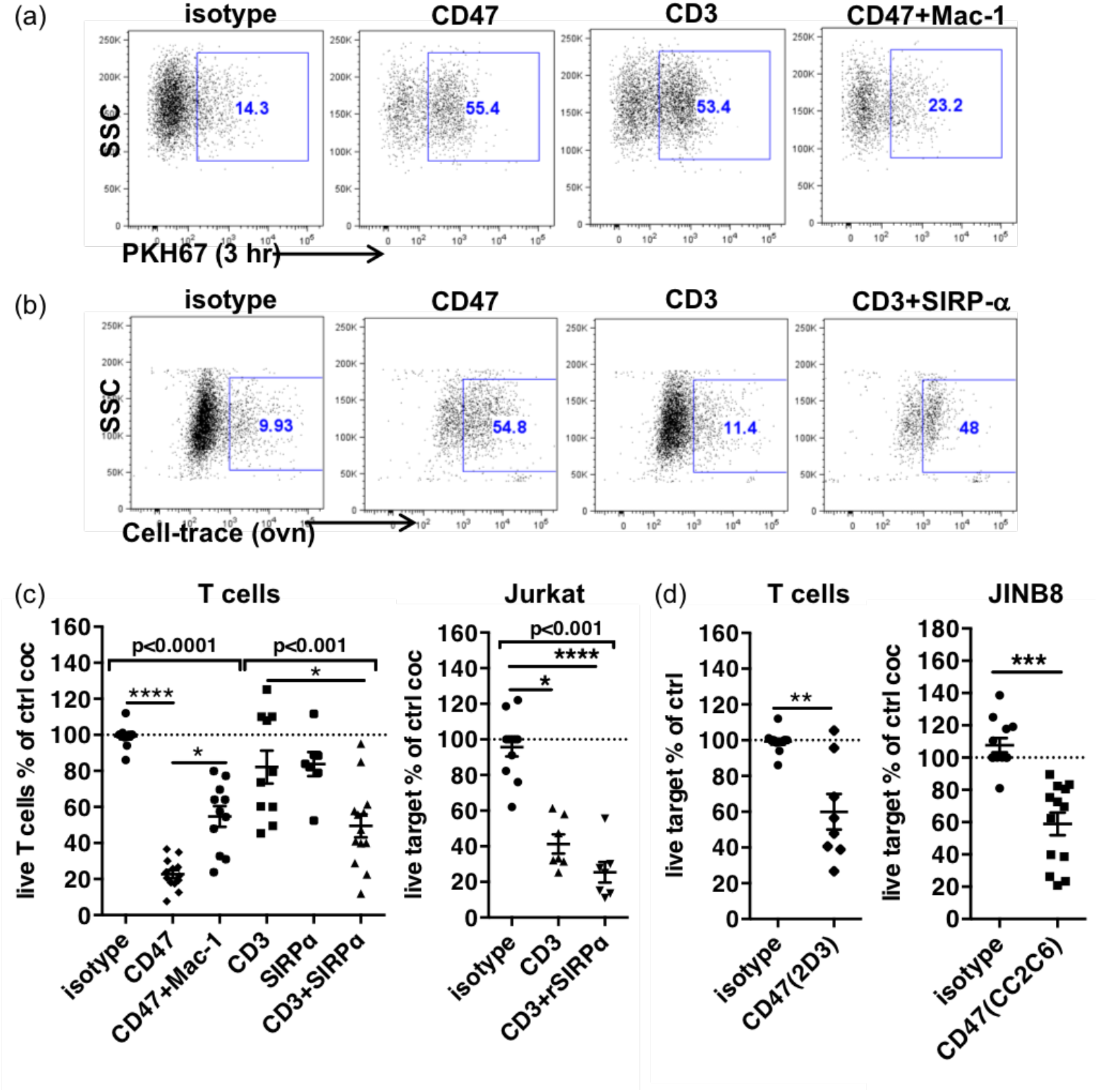
Trogocytosis and cytotoxicity in PMNs’ ADCC. (a) Expression of the PKH67 T cells membrane dye in PMNs after a 3-hours coculture in the presence of indicated mAbs. (b) Uptake of CellTrace stained T cells components in PMNs after overnight coculture in the presence of mAbs. For (a)(b) PMNs are gated after exclusion of doublets and dead cells. (c)(d) Cytotoxicity of PMNs to targets indicated on top of graphs, evaluated by the percentages of live targets compared to control after overnight coculture. N=6-12 different donors, ratios of PMNs to target=1-3. CD3: anti-CD3 mAbs; CD47: anti-CD47 mAbs clone CC2C6 or 2D3 as indicated; Mac-1: anti-CD11b+anti-CD18 mAbs. SIRPα: anti-SIRPα mAbs. rSIRPα: recombinant SIRPα protein. Median and SEM. *P*-values from Kruskall-Wallis test indicated on top of groups, *P*-values from Dunn’s multiple comparison post-test on top of pairs: *, *P*<0.05; **, *P*<0.01; ***, *P*<0.001.

These results suggested that this immediate trogocytosis was not controlled by CD47-SIRPα signaling but via FcR activation. We investigated trogocytosis in different experimental conditions using CellTrace Violet labelled T cells incubated overnight with PMNs (as in Matlung) (Figure 2-b, Figure S3-a). The uptake of CellTrace in PMNs by anti-CD3 mAbs was not different from control (11% vs 9.9% resp.) but blockade of CD47-SIRPα interaction with anti-SIRPα antibodies increased it to a level similar to anti-CD47 mAbs (48% and 54.8%, resp.) consistent to a control of SIRPα.

Next we compared the cytotoxicity induced in overnight cocultures of PMNs with T cells. At ratios ranging from 1-3:1, anti-CD47 mAbs-ADCC decreased T cells viability of a factor of 5 (22% survival) (Figure 2-c, Figure S2-b). Anti-Mac-1 mAbs only partially restored T cells survival (54% vs 22%) while fully inhibited trogocytosis, suggesting that adhesion-mediated ADCC was involved in cytotoxicity but does not account for the complete effect. Consistent with a trogocytosis mechanism, PMNs’ cytotoxicity induced by anti-CD3 mAbs was weak but became significant when anti-SIRPα mAbs were added to block the “don’t eat me” pathway (82% vs 50% survival, resp.). Similarly, blockade of SIRPα with a recombinant SIRPα protein (rSIRPα) (as in (33)) increased cytotoxicity of PMNs to Jurkat T cells opsonized with anti-CD3 mAbs by a factor of 1.6-fold (41% vs 25%, resp.) (Figure 2-c). In the two systems however, blockade of SIRPα did not result in the same level of cytotoxicity induced by anti-CD47 mAbs. Although the direct effect of anti-CD47 mAbs on T cells must slightly contribute to the whole effect in the presence of PMNs (see Figure 1-b), the combination of anti-CD3 *plus* anti-SIRPα mAbs seemed unable to induce similar level of PMN-mediated cytotoxicity despite inducing similar trogocytosis.

These results suggested that another mechanism than ADCC *plus* blockade of the engagement of SIRPα was involved in the cytotoxicity induced by anti-CD47 mAbs. To demonstrate more directly this hypothesis, we used an anti-CD47 mAb targeting an epitope outside the interaction site with SIRPα (2D3) to induce ADCC against T cells and a CD47-deficient Jurkat T cell line as target for PMNs in the presence of anti-CD47 mAb CC2C6 (Figure 2-d, Figure S1-b, Figure S2-c). Although weak effect was expected, PMNs displayed a significant cytotoxicity in the two systems demonstrating the contribution of additional mechanisms likely resulting from the CD47 triggering on PMNs in the strong cytotoxicity.

### Triggering of CD47 on PMNs induced a strong production of ROS

To address the role of PMN CD47 engagement vs. T cell CD47 engagement in cytotoxicity, we pre-treated PMNs or T cells for 30 min with the anti-CD47 mAb CC2C6 then washed the antibody before co-culture with T cells. Only pre-treatment of PMNs resulted in a significant cytotoxicity to T cells, however weaker than that obtained when the antibody was present throughout the co-culture (57% and 18%, resp., Figure 3-a).

**Figure 3.**
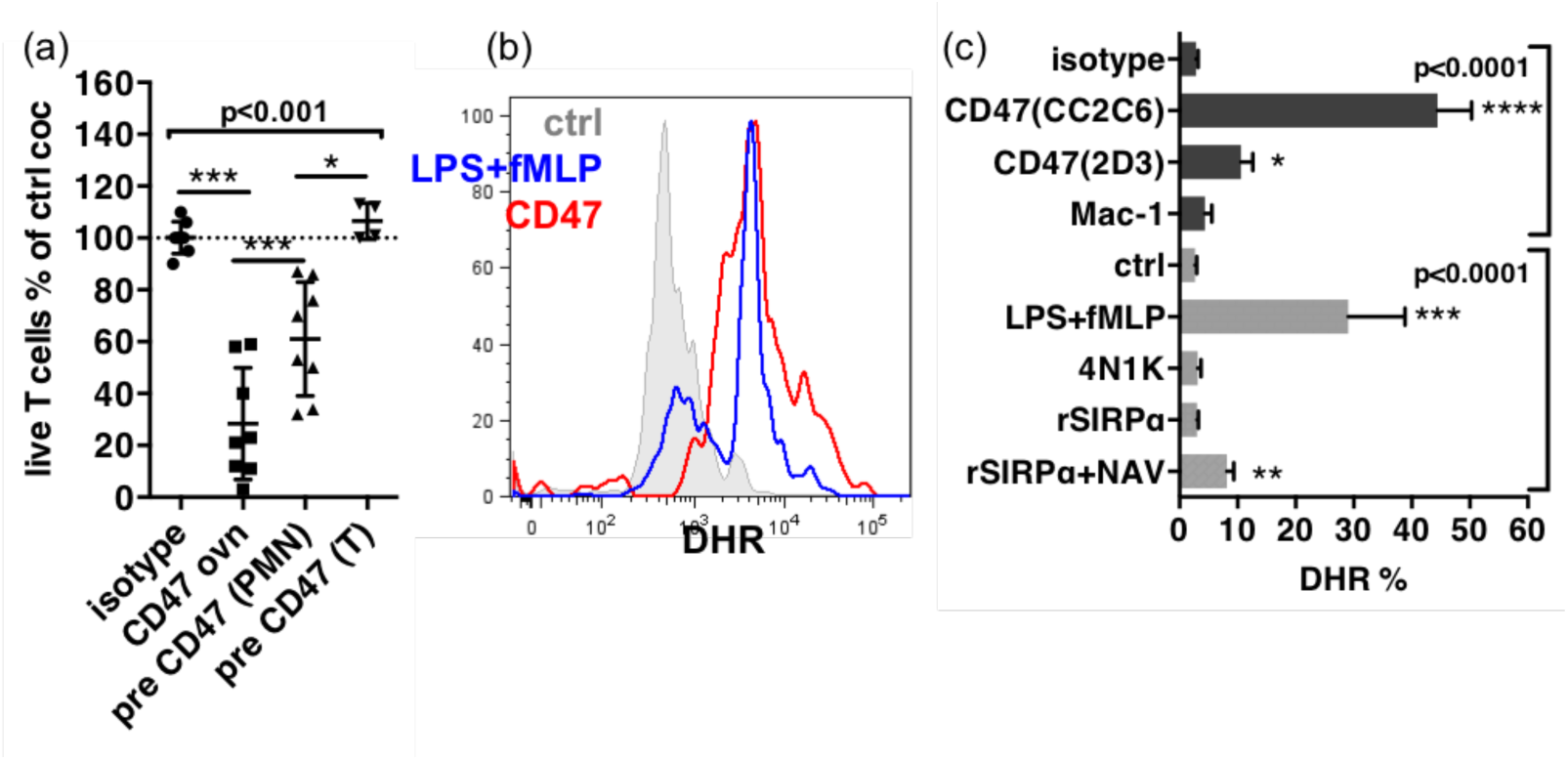
Induction of production of ROS in PMNs. (a) Effect of pre-treatment of cells by the anti-CD47 mAb clone CC2C6 on cytotoxicity of PMNs to T cells. Cytotoxicity is represented by the percentages of live T cells compared to control coculture (ctrl coc). Pre-CD47: pre-treatment 30 min at 4°C. CD47 ovn: anti-CD47 mAb during overnight culture, n=8. (b) Histograms of DHR staining in PMNs after exclusion of dead cells and doublets after 1 hr stimulation. (c) Percentages of expression of DHR in PMNs after 1 hr stimulation. N=15-25 for antibodies, n=5 for recombinants proteins or peptides. Gates are set on unstimulated stained cells.

Since PMN are potent producers of ROS we investigated whether PMNs stimulated by anti-CD47 mAbs produced ROS using DHR staining. LPS *plus* fMLP were used as a positive control for ROS induction, isotype as a negative control. Anti-CD47 mAb CC2C6 induced a rapid increase of DHR fluorescence in PMNs similar to LPS *plus* fMLP (Figure 3-b, Figure S4-a). This result suggested that antibody interaction with CD47 was responsible for the activation of NADPH oxidase, resulting in production of ROS. This hypothesis was consistent with the known lateral association of CD47 with GPCR involved in NADPH oxidase activation (19). Consistently, the 2D3 mAb also induced ROS (Figure 3-c, Figure S4-a).

Likewise, weaker binding recombinant ligands of CD47 like the 4N1K peptide (34) and recombinant SIRPα (29) failed to induce ROS except when rSIRPα was multimerized with neutravidin to increase significantly (however weakly) DHR, suggesting that avidity or membrane clustering might be important factors of direct ROS induction through CD47 (Figure 3-c). However, the weaker activation of ROS production by direct triggering than the SIRPα-blocking CC2C6 might also suggest that CD47 engagement alone does not explain the whole effect of the latter.

### Production of ROS is activated in PMNs’ ADCC and tightly controlled by SIRPα

Although results with isotype and anti-Mac-1 mAbs demonstrated that FcR stimulation was not involved in induction of ROS, anti-CD47 mAbs CC2C6 have nevertheless an intact Fc domain and unexpectedly, anti-SIRPα mAbs also induced ROS (Figure 4-a, Figure S4-a). This result prompted us to consider the hypothesis of the activation of NADPH oxidase by a reciprocal FcR activation in PMNs, or by a trimolecular complex between known as the scorpion effect (35) previously observed with anti-SIRPα antibodies on macrophage (36). ROS were activated only by anti-CD47 or anti-SIRPα mAbs but not by anti-Mac-1 mAbs, i.e. only in the context of the simultaneous blockade of SIRPα. This hypothesis implied thus that if NADPH oxidase was activated through FcR stimulation in PMNs, it was simultaneously and strictly controlled by the engagement of SIRPα a feature not yet described in PMN.

**Figure 4.**
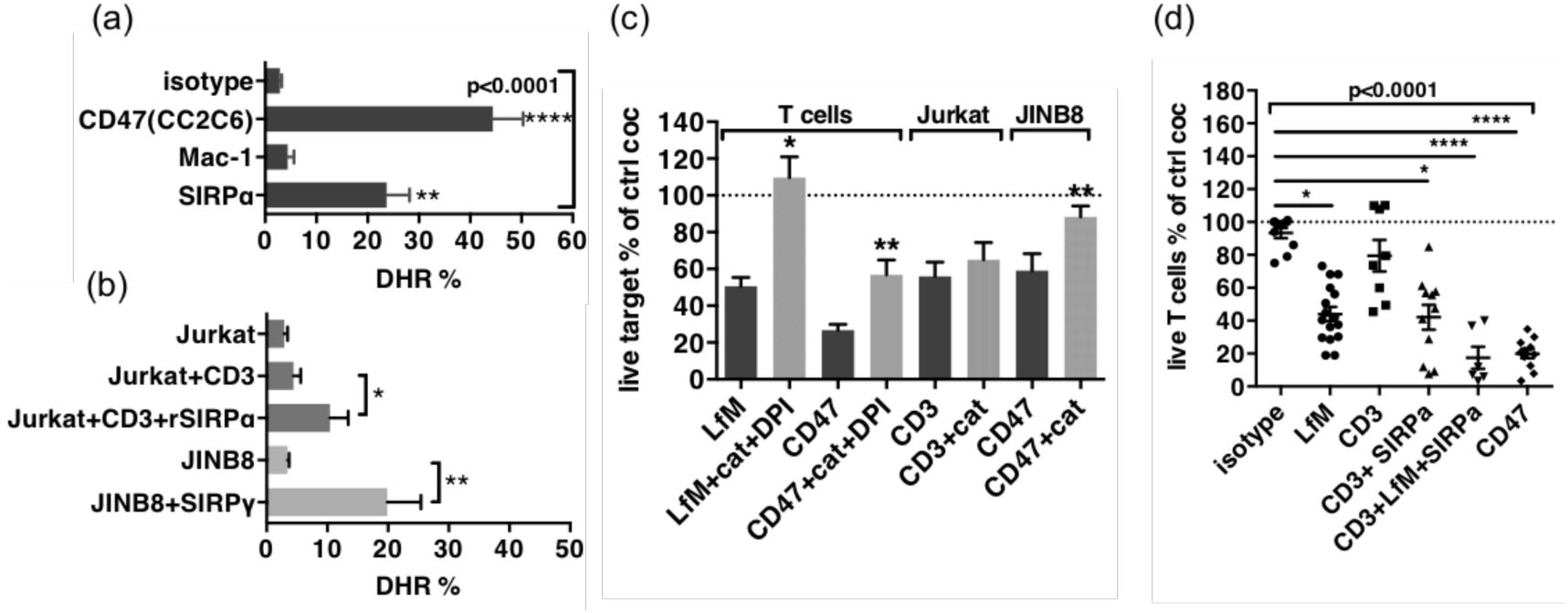
Production of ROS is stimulated in ADCC and contributes to cytotoxicity. (a) Stimulation of ROS in PMNs by mAbs to PMNs. (b) Stimulation of ROS in PMNs by cocultures with targets and mAbs. For (a) and (b), Median and SEM of percentages of expression of DHR in PMNs after 1 hr stimulation. For(a), n=6-25 different donors. *P*-values from Kruskall-Wallis test indicated on top, *P*-values from Dunn’s multiple comparison post-test on top of pairs: **, *P*<0.01; ***, *P*<0.0001. For (b) n=6-10. *P*-values of Wilcoxon matched pair-test: * *P* < 0.05; ** *P*<0.01. (c) Inhibition of cytotoxicity by catalase (cat) and DPI, n=6-25. Median and SEM, *P*-values of Wilcoxon matched pair-test: * *P* < 0.05; ** *P*<0.01. (d) Reconstitution of the cytotoxicity induced by anti-CD47 mAb CC2C6 (CD47) using anti-CD3 (CD3) + anti-SIRPα (SIRPα) + LPS + fMLP (LfM), n=6-14. Median and SEM, *P*-values from Kruskall-Wallis test indicated on top, *P*-values from Dunn’s multiple comparison post-test on top of pairs: *, *P*<0.05; **, *P*<0.01; ***, *P*<0.0001.

To demonstrate this hypothesis, we investigated intracellular induction of ROS in short co-cultures of PMNs with targets (Figure 4-a, Figure S4-b). Coculture with Jurkat opsonized by anti-CD3 mAbs did not significantly increase DHR in PMNs (5% and 4%, resp.), while the presence of rSIRPα (to block interaction with SIRPα) significantly increase DHR to 10%. Similarly, coculture of PMNs with CD47-deficient target cells (JINB8) opsonized by anti-SIRPγ mAbs significantly increased DHR in PMNs to 20%. These results suggested that stimulation of PMNs through FcR induced ROS but only when PMN-SIRPα was not engaged by target cell CD47, either by blockade of CD47 with rSIRPα or by the absence of CD47 on target.

We evaluated the contribution of ROS in cytotoxicity induced by PMNs’ ADCC on T cells using catalase, a H_2_O_2_ scavenger *plus* diphenyleneiodonium chloride (DPI), an NADPH oxidase inhibitor, to counteract ROS effects during coculture and stimulation. LPS *plus* fMLP was used to induce production of ROS by a pathway independent of ADCC. The resulting cytotoxicity on T cells was significant but weaker than anti-CD47 mAbs (50% vs 26% survival resp., Figure 4-c) and fully blocked by catalase plus DPI. The cytotoxicity induced by anti-CD47 mAbs was significantly but partially reduced to 56% while the significant cytotoxicity induced by anti-CD3 mAbs on Jurkat T cells in absence of blockade of SIRPα was not modified (Figure 2-c, Figure 4-c). These results suggested that ROS produced during ADCC in the context of blockade of SIRPα contributed to the cytotoxicity of PMNs. Finally, we verified our hypothesis by using together anti-CD3 mAbs to induce ADCC, anti-SIRPα mAbs to block the inhibitory checkpoint and LPS *plus* fMLP to induce ROS. This combination reached the levels of cytotoxicity induced by anti-CD47 mAbs, suggesting the important contribution of ROS in the process (Figure 4-d, Figure S3-b).

## Discussion

In the context of the toxicities generated by blockade of the SIRPα-CD47 checkpoint in cancer therapies, we sought to address the role of PMN in the lymphopenia observed with anti-CD47 antibody treatment (25). We found that the strong PMN-mediated cytotoxicity induced by the anti-CD47 mAbs CC2C6 was sustained not only by trogocytosis but also by a strong respiratory burst, the cooperation of both resulting in a uniquely efficient killing of T cells in a 1:1 effector cell to target cell ratio.

The anti-CD47 mAb clone CC2C6 can induce a weak T cells death through direct triggering (31). In coculture with PMNs, this antibody can stimulate FcR on PMNs and simultaneously blocks CD47 interaction with SIRPα resulting in potent ADCC (4, 37). In this regard, we found that PMNs killed leukemic B cells in the presence of anti-CD47 antibodies regardless of the presence of the well-known ADCC antibody Rituximab. This result suggested that anti-CD47 mAbs alone were sufficient for a cytotoxic response and explained why PMNs also killed T cells. However, the reconstitution of this scenario using a T cell opsonizing antibody (anti-CD3) that induces ADCC *plus* anti-SIRPα mAbs or recombinant SIRPα protein to block the “don’t eat me” checkpoint did not result in such strong killing suggesting that another mechanism must contribute to the cytotoxicity induced by anti-CD47 mAbs.

The most likely hypothesis was that this additional mechanism was triggered by the interaction of the antibody with CD47 on PMNs. This was addressed by measuring the anti-CD47-induced cytotoxicity of PMN cells in coculture with CD47-deficient T cells (Jurkat T cell clone JINB8). Though less toxicity was observed when compared to CD47^+^ T cells, we still observed a significant increase in killing as compared to control treatment. These observations suggest that targeting CD47 on PMN is sufficient to activate cytotoxic mechanisms.

We investigated whether respiratory burst, the most common mechanism used by PMNs to efficiently kill their targets in a short time, was involved. Not surprisingly ROS contributed to the strong killing of T cells by PMNs induced by anti-CD47 mAbs. This was demonstrated by i) direct induction of ROS in PMNs by stimulation with anti-CD47 mAbs, ii) partial blockade of cytotoxicity by catalase and DPI, iii) reconstitution of equivalent cytotoxicity by induction of ADCC with anti-CD3 mAbs *plus* blockade of SIRPα with antibodies *plus* stimulation of ROS with LPS and fMLP.

The next question was to determine how anti-CD47 mAbs stimulated NADPH oxidase in PMNs. Direct activation of CD47 was suggested by the significant cytotoxicity observed during pre-treatment of PMNs with CC2C6 as well as the induction of intracellular ROS by direct treatment of PMNs with CC2C6 and 2D3, an anti-CD47 mAb that targets an epitope outside the site of interaction with SIRPα (37). This hypothesis was moreover supported by the known lateral association of CD47 with GPCR, themselves involved in NADPH oxidase activation (14, 19). A higher affinity multimer of the recombinant SIRPα protein obtained with neutravidin (KD = 16 nM, (33)) reproduced a significant but weak induction of ROS while other soluble ligands failed, suggesting that affinity or CD47 clustering might be critical. These data support direct linking of CD47 as a pathway of activation of NADPH oxidase, as recently suggested on T lymphomas Jurkat cells (38) However, anti-SIRPα mAbs also induced production of ROS in PMNs although SIRPα is known primarily as an inhibitory receptor (5). This contradictory result prompted us to consider an alternative mechanism that could result from stimulation of FcR either by reciprocal interactions between PMNs or by formation of trimolecular complexes with FcR on each cell (35). Induction of ROS occurred only with whole antibodies that simultaneously to stimulate FcR blocked SIRPα but not with those that could not (like anti-Mac-1 mAbs). Thus, this proposed mechanism would imply that the activation of NADPH oxidase through FcR is controlled by engagement of SIRPα. Considering the potential cytotoxicity of ROS, this tight control is not surprising. Support for this hypothesis was demonstrated by the induction of ROS in PMNs co-cultivated with opsonized targets either deficient for CD47 or where engagement of SIRPα by CD47 was blocked by recombinant SIRPα protein, that fails itself to stimulate ROS. Such control of SIRPα on NADPH oxidase activation was previously shown during myeloid cells differentiation by the restriction of the expression of the gp91^Phox^ subunit of NADPH oxidase (15). The mechanism may be different in mature PMNs considering the short time of response to stimulation and the basal expression of gp91^Phox^ in mature cells.

Although our results confirmed the activation of NADPH oxidase in FcR stimulation previously proposed by van Spriel *et al*. (7), recent work based on PMNs from NADPH oxidase deficient patients claimed that trogoptosis, a mechanism resulting from lytic processes induced by trogocytosis, was solely responsible for ADCC of PMNs (10). Our results are not contradictory with this statement: ADCC without blockade of SIRPα can only rely on trogoptosis and is efficient provided that high ratios of effector to target are used. Ratios of 50:1 are used to kill 30% of SKBR3 breast cancer cells opsonized with Trastuzumab (10, 12) and killing increases of only 2-fold on a CD47-deficient target. On the contrary, trogocytosis at low ratios of PMN to CLL-B cells opsonized with anti-CD20 mAbs does not induce significant death (11) unless SIRPα is blocked allowing activation of NADPH oxidase and strong killing, as we show here for ratios of 1:1. So cytotoxicity can either result from trogocytosis alone or ROS alone but when the two mechanisms cooperate a single PMN becomes a potent killer. It is tempting to speculate that the spill of ROS into the intracellular milieu of targets through membrane holes created by trogocytosis is the mechanism of optimal function of PMNs. This relationship between number of PMNs focusing on a target and underlying mechanisms of killing opens new perspectives on the fundamental biology of PMNs where SIRPa-CD47 appears as a key regulator of PMN capacity to differentiate between physiological trogocytosis and cytotoxicity by unleashing trogocytosis and ROS.

The new information brought by our work on the role of NADPH oxidase activation in ADCC of PMN and its control by SIRPα might help to design new agents aimed to enhance myeloid cells function in the treatment of cancer while limiting adverse effects on healthy cells, in particular those generated by PMNs. Targeting CD47 with FcR-stimulating antibodies is less than ideal as it is responsible of collateral killing of non-tumor cells bearing CD47. Targeting CD47 with synthetic agents would induce the full destructive capacity of PMN by activation of NADPH oxidase through direct linking in addition to blockade of SIRPα. However, although ROS are expected to be preferentially released into tumor cells carrying therapeutic antibodies subjected to trogocytosis, collateral damage to non-opsonized cells due to release of ROS in the milieu are not negligible, especially for T cells that appear highly sensitive in our studies. Therefore, from the perspective of PMN activity, targeting CD47 using a non-ADCC inducing agent or targeting SIRPα appear to be approaches that can both maximize tumor targeting/killing while limiting collateral damage to non-tumor and potentially tumor-hostile immune cells. Indeed, these strategies in syngeneic mouse tumor models stimulated both innate (macrophage, dendritic cells, and PMN) as well as innate (cytotoxic T cell) responses (39, 40). This is evident in their ability to enhance T cell directed therapies (anti-PD-1/PD-L1) when used in combination. This would also ideally target PMN already in the tumor microenvironment as modulation of SIRPα signaling is known to impede PMN transmigration (6).

## Acknowledgments

The authors are grateful to Manon Richaud and Françoise Mallet for assistance with the use of the Cytometry Platform of the CRCM. TWM also wishes to thank the programmatic staff at the Preclinical Therapeutics Grants Branch of the NCI for their support of this research program.

## Notes

Additional information: **Financial Support:** this work was supported by the National Cancer Institute (1U01CA218259, R37 CA218259; TWM), the Fondation ARC for Cancer Research (COVID202001312), the team “Immunity and Cancer” was labeled by the Fondation pour la Recherche Médicale “Equipe FRM DEQ20180339209”. DO is Senior Scholar of the Institut Universitaire de France.

### Competing Interest Statement

DO is a cofounder of Imcheck Therapeutics, Alderaan Biotechnology and Emergence Therapeutics. TWM is a co-owner of Paradigm Shift Therapeutics. The other authors declare no competing financial interests.

