## Supplemental Figures for "CD47-SIRPα controls ADCC killing of primary T cells by PMN through a combination of trogocytosis and NADPH oxidase activation"

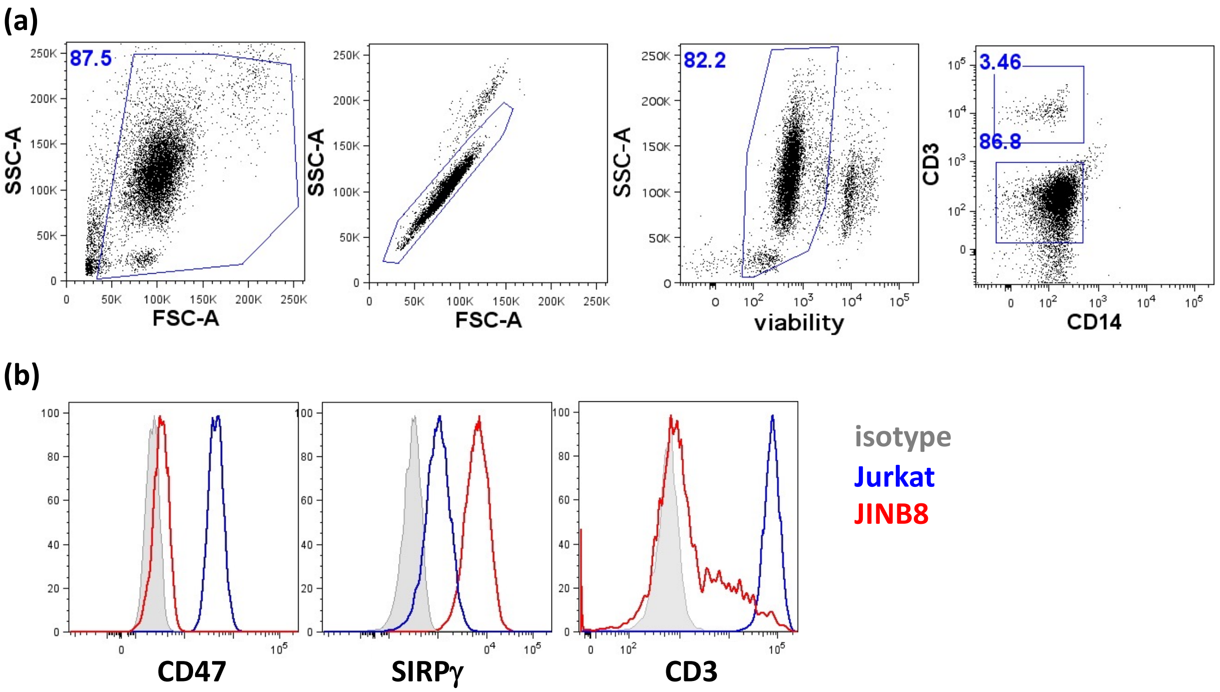


**Fig S1. Characterization of cells.** (a) Purity of PMNs' preparations. Dot plots shows sequential gating of PMNs after separation from blood. From left to right, exclusion of debris, of doublets, of dead cells and analysis of contamination with monocytes $CD14^+$ and T cells $CD3^+$. (b) Expression markers on Jurkat (JA16 clone) and Jurkat CD47-deficient cells (JINB8 clone).

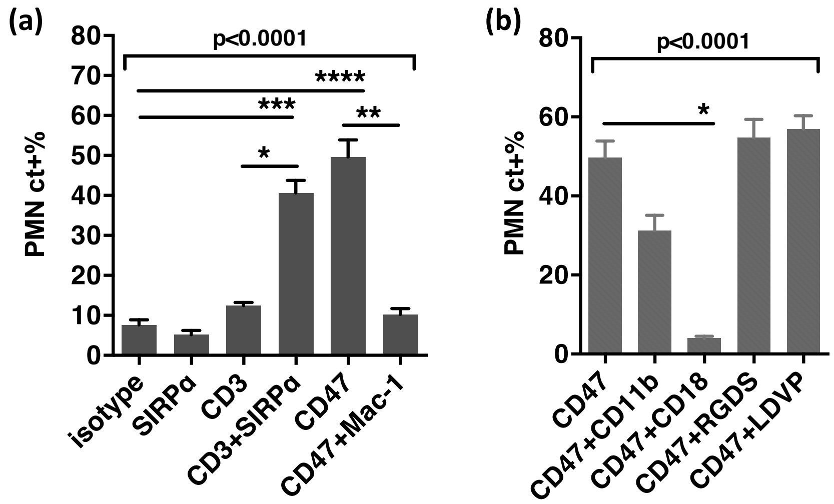


**Fig. S2. Trogocytosis of PMNs.** (a) Statistical analysis of trogocytosis of PMNs to cell-trace stained T cells in indicated conditions. Trogocytosis is represented by percentages of cell-trace in PMNs, gated after exclusion of doublets, dead cells and cell-trace$^{+}$CD3$^{+}$ targets. (b) Inhibition of trogocytosis of PMNs to T cells by blockade of adhesion with mAbs (CD11b, CD18) or peptides (RGDS, LDVP), n=6-12 experiments with different PMN's donors. *P*-values from Kruskall-Wallis test is indicated on top of group, *P*-values from Dunn's multiple comparison post-test on top of pairs :*, $P<0.05$..

**(a) ADCC of PMNs to Raji lymphoma B-cells**

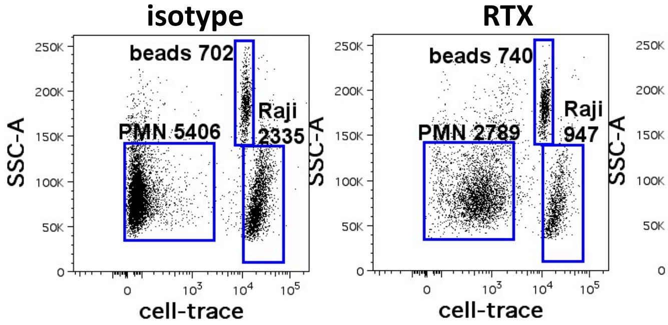


**(b) ADCC of PMNs to T cells**

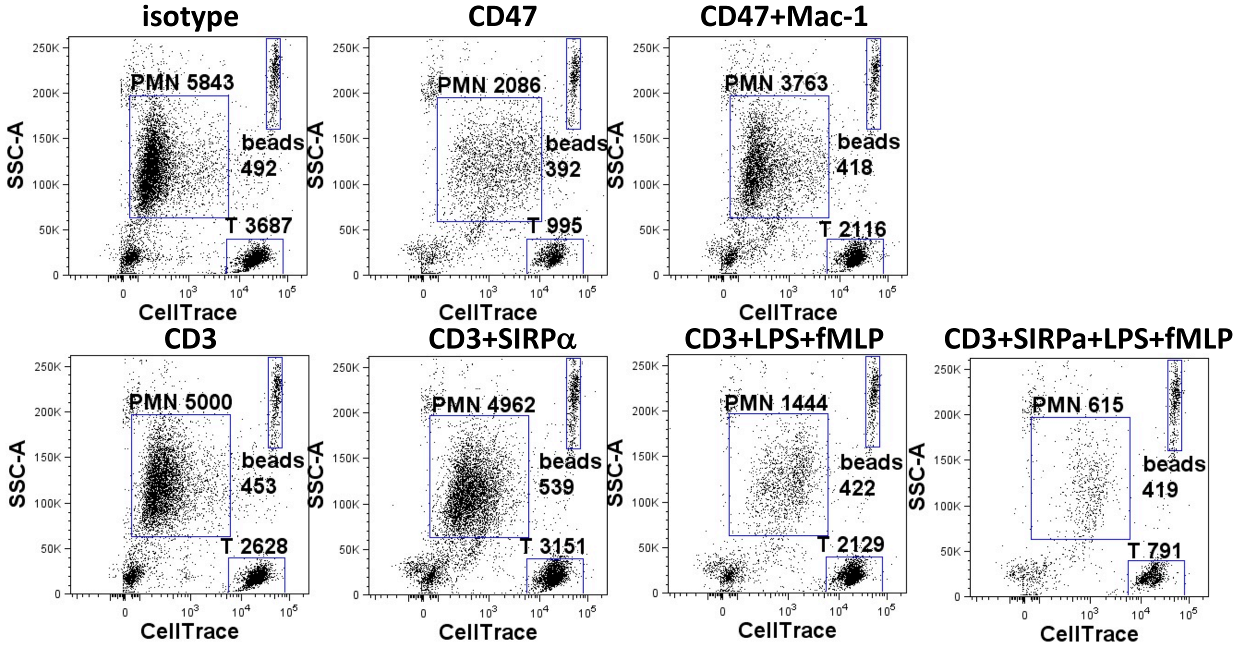


**(c) Induction of PMNs' cytotoxicity to JINB8 by anti-CD47 mAbs**

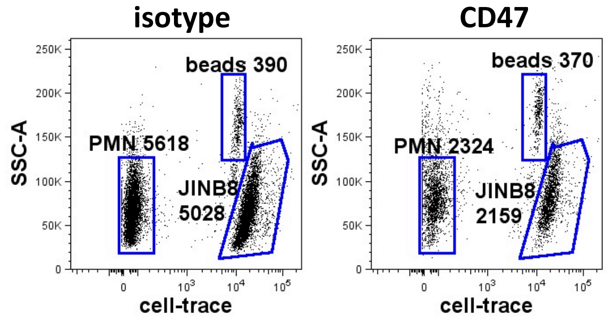


**Fig S3. Cytotoxicity assays.** Analysis of survival of targets after overnight co-cultures with PMNs. (a) Raji B-lymphoma cells (ratio of PMNs to targets=2 at T0); (b) T cells, (ratio of PMNs to T cells=0.84 at T0); (c) JINB8 (ratio of PMNs to JINb8=0.8 at T0). RTX: Rituximab; CD47: anti-CD47 mAbs; Mac-1: anti-CD11b plus anti-CD18 mAbs; SIRPα: anti-SIRPα mAbs; CD3: anti-CD3 mAbs. Dot plots show cell-trace/SSC-A after exclusion of doublets and dead cells. Gates and counts of beads, PMNs and targets.

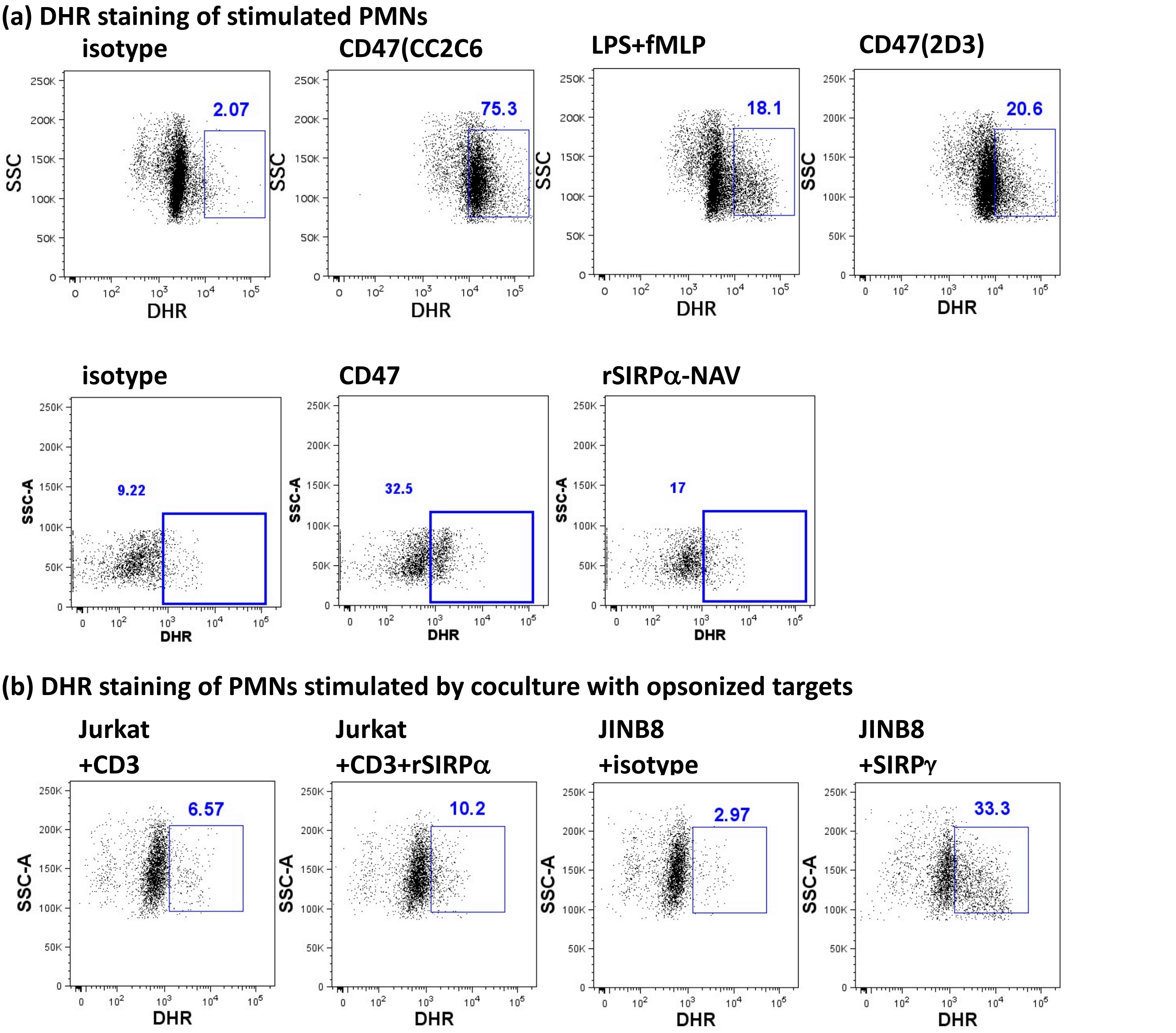


**Fig S4. Induction of ROS in PMNs.** (a) and (b) Dot plots show DHR staining of PMNs gated after exclusion of doublets, dead cells and cell-trace stained targets as $SSC^{hi}$ cells in indicated stimulation conditions. CD47: anti-CD47 mAbs; CD3: anti-CD3 mAbs ; SIRPγ: anti-SIRPγ mAbs; rSIRPα: recombinant SIRPα; NAV:neutravidin. Percentage of DHR expressing cells within PMNs is indicated in each graph.
